# Altered TGFB1 regulated pathways promote accelerated tendon healing in the superhealer MRL/MpJ mouse

**DOI:** 10.1101/2021.02.08.430308

**Authors:** Jacob G. Kallenbach, Margaret A. T. Freeberg, David Abplanalp, Jacquelyn A. Myers, John M. Ashton, Alayna Loiselle, Mark R. Buckley, Andre J. van Wijnen, Hani A. Awad

## Abstract

To better understand the molecular mechanisms of tendon healing, we investigated the Murphy Roth’s Large (MRL) mouse, which is considered a model of mammalian tissue regeneration. We show that compared to C57Bl/6J (C57) mice, injured MRL tendons have reduced fibrotic adhesions and cellular proliferation, with accelerated improvements in biomechanical properties. Transcriptional analysis of biological drivers showed positive enrichment of TGFB1 in both C57 and MRL healing tendons. However, only MRL tendons exhibited downstream transcriptional effects of cell cycle regulatory genes, with negative enrichment of the cell senescence-related regulators, compared to the positively-enriched inflammatory and ECM organization pathways in the C57 tendons. Serum cytokine analysis revealed decreased levels of circulating senescence-associated circulatory proteins (SASP) in response to injury in the MRL mice compared to the C57 mice. These data collectively demonstrate altered TGFB1 regulated inflammatory, fibrosis, and cell cycle pathways in flexor tendon repair.

## Introduction

Flexor tendons transmit the intricate muscle forces required to move the digits freely. These hierarchically structured and tightly aligned collagen-rich connective tissues are sparsely populated and maintained by tenocytes (Nichols et al., 2019), which are fibroblast-like cells that express phenotypic transcriptional factors including scleraxis (Scx) (Best and Loiselle, 2019, Pryce et al., 2007, Pryce et al., 2009, Schweitzer et al., 2001), mohawk (Mkx) (Berthet et al., 2013, Ito et al., 2010, Liu et al., 2010), the tendon matrix protein of collagen type I (Col1), and non-collagenous proteins such as the glycoprotein tenomodulin (Tnmd) (Alberton et al., 2015, Dex et al., 2017, Lin et al., 2017, Qi et al., 2012). Injuries to flexor tendons, particularly in zone II of the hand, are susceptible to debilitating fibrotic adhesions, which compromise tendon gliding, strength, and range of motion, which is often attributed to the tendon’s sparse cellularity (Whalen, 1951) and vascularity (Tempfer and Traweger, 2015). Upon injury, tendon healing proceeds in three canonical and overlapping phases: 1) an inflammation phase, 2) a proliferative, reparative phase, and 3) a remodeling phase (Nichols et al., 2019, Andarawis-Puri and Flatow, 2018, Thomopoulos et al., 2015). Immediately following tendon injury, acute cytokine and chemokine production signals circulating and peripheral inflammatory cells, predominantly leukocytes, to the hematoma forming at the injury site. The proliferative, reparative phase is driven by intrinsically and extrinsically derived fibroblasts, including the tenocytes, fibroblasts from the loose connective tissues enveloping the tendon (epitenon or paratenon) and its collagenous fascicles (endotenon) and perivascular cells (Dyment and Galloway, 2015). These fibroblasts scaffold a fibrovascular provisional matrix (granulation tissue) rich with collagen type III to partially restore tissue continuity. The remodeling phase, which is typically protracted, gradually replaces the provisional matrix with native tissue ECM components that are restructured to restore uninjured tissue mechanical properties, mediated by enzymatic activity of matrix metalloproteinases (MMPs). Time-dependent activation of Scx-expressing cells within the injury microenvironment is thought to constitute regenerative tendon healing, whereas the activation of α-SMA^+^ myofibroblasts precipitates scar-mediated healing (Kaji et al., 2020, Dyment et al., 2013, Best and Loiselle, 2019). Fibrotic scar persistence is thought to occur from an inadequate regenerative response from the native Scx-expressing cells, chronic presence of inflammatory cells and signals, heightened myofibroblast activity, and aberrant MMP remodeling (Nichols et al., 2019). Despite recent advances elucidating the cellular and molecular mechanisms of tendon healing, gaps in knowledge remain and have hindered the identification of novel therapeutic candidates that could prevent fibrosis and stimulate regenerative tendon healing.

Multifarious studies have identified the Murphy Roth’s Large (MRL/MpJ) mouse as a promising mouse model of mammalian tissue regeneration due to its ability to restore native-like tissue structures and properties within the ear (Clark et al., 1998), skin (Beare et al., 2006), skeletal muscle (Heydemann et al., 2012, Mull et al., 2014, Sinha et al., 2019, Tseng et al., 2019), articular cartilage (Fitzgerald, 2017, Fitzgerald et al., 2008, Leonard et al., 2015, Mak et al., 2016, Rai and Sandell, 2014, Ward et al., 2008), and tendon (George et al., 2020, Lalley et al., 2015, Paredes et al., 2020, Paredes et al., 2018, Sereysky et al., 2013). Originally hypothesized that the immunomodulated systemic environment in MRL mice drove these regenerative phenotypes (Fitzgerald, 2017, Fitzgerald et al., 2008), studies show the necessity of blood flow or vasculature to recapitulate native tissue structures such as skin (Davis et al., 2005), while more avascular tissue injuries such as cartilage demonstrate minimal improvements (Fitzgerald, 2017, Fitzgerald et al., 2008). Even though uninjured tendon is hypovascular, a recent patellar tendon defect study removed the central third of the patellar tendon from MRL and C57BL/6J (C57) mice and demonstrated that MRL patellar tendons recapitulate near native biomechanical properties (Lalley et al., 2015). More recent studies injured the patellar tendon with a 0.75mm tendon mid-substance biopsy punch and again demonstrated that MRL patellar tendons heal to near native biomechanical properties with an attenuated inflammatory response, heightened MMP activity, and improved ECM alignment (George et al., 2020, Paredes et al., 2020, Paredes et al., 2018). While the MRL enhanced regenerative potential within musculoskeletal tissues may depend on tissue-specificity and injury severity, recent transcriptomic (Podolak-Popinigis et al., 2015, Sebastian et al., 2018), proteomic (Caldwell et al., 2008), and metabolomic (Tseng et al., 2019, Podolak-Popinigis et al., 2015) techniques point to novel mechanistic insights. Altogether, the prevailing evidence suggests that the MRL tendon tissue healing response is improved, but the specific cellular and transcriptomic mechanisms of these biological effects remain to be elucidated.

In this study, we report histological and biomechanical data indicative of accelerated tendon repair in the MRL superhealer model in comparison to standard C57 mice. Our transcriptional analysis suggests that these biological effects are related to altered TGFB1 regulated gene sets negatively enriching cell cycle and senescence transcriptional pathways in the MRL tendons, and positively enriching inflammatory and ECM organization pathways in the C57 tendons.

## Results

### MRL tendons heal with reduced adhesions and improved biomechanical properties

We first sought to compare the extent of adhesion formation and repair strength of healing flexor tendons in the C57 and MRL mice. The healing of partially lacerated deep digital flexor tendons (DDFT) was qualitatively assessed using hematoxylin-stained axial tendon sections. In general, the MRL tendons appeared to respond to the injury with a reduced cellular response compared to the C57 tendons. Specifically, there were no gross morphological differences in the uninjured (Fig. 1 A,E), where the tendon bundles were sparsely populated by cells, and the tendons were encased with a very thin synovial sheath (black arrows in Fig. 1 *a,e*). Fourteen days post injury (dpi), injured tendons in both mouse strains experienced substantial hyperplasia in the surrounding synovial sheath (black arrows, Fig. 1 *b,f*), while no significant changes in cell number in the tendon proper were generally observed (yellow arrows Fig. 1 B,F). At 28 dpi, substantial increases in cell density in both the tendon bundle and the peritendinous adhesions were observed in the C57 mice but not the MRL mice (yellow arrows, Fig. 1 C,G). At this stage, the synovial hyperplasia began to resolve in the MRL mice but persisted in the C57 mice (black arrows, Fig. 1 *c,g*). By 56 dpi, the injured MRL tendons appeared to have restored uninjured morphology where the tendon bundle was sparsely cellular and encased by a thin synovial membrane (Fig. 1H,*h*). In contrast, while the cellularity of the C57 tendon bundle decreased, the surrounding synovial sheath still demonstrated a hyperplasia morphology at 56 dpi (Fig. 1D,*d*). Quantification of the tendon perimeter adhered to the surrounding sheath and tissues (Fig. 1I) and cellular density (Fig. 1J) in the tendon bundle demonstrated significant increases in adhesions and cellularity in the C57 tendons at 14 dpi and 28 dpi, respectively, compared to the MRL tendons.

**Figure 1.**
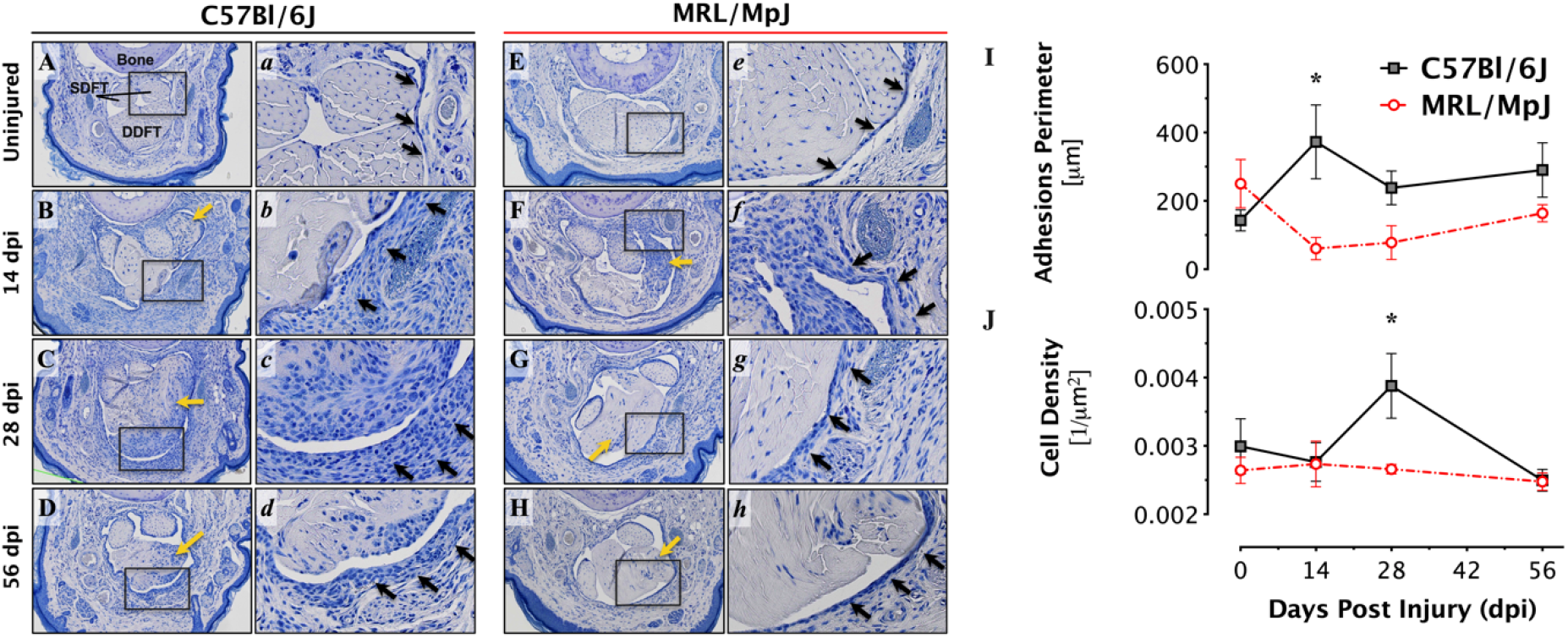
MRL flexor tendons heal with reduced peritendinous adhesions. Representative (2X) micrographs (A-H) of hematoxylin-stained cross-sections of the digits of C57 and MRL mice before and after zone II injury up to 56 days post injury (dpi). High magnification (20X) regions of interest (*a*-*h*) are representative of five levels analyzed per tendon. Black arrows identify the thin synovial sheath in uninjured tendons, and the hyperplasia that ensues immediately after injury in both strains. The synovial sheath hyperplasia appears to persist in the C57 tendons up to 56 dpi, but resolves in MRL tendons by 28 dpi. Yellow arrows indicate regions of increased cellularity in the injured tendon and peritendinous tissues evident by the punctate, intense hematoxylin staining. Histomorphometric quantification of the healing tendon shows reduced adhesions (I) and tendon hyperplasia (J) in the MRL mouse. The length of the tendon perimeter adhered to the subcutaneous tissue and cell density are significantly reduced at 14- and 28- dpi in the MRL, respectively (n=4).

To test whether the changes in cell density over time and between strains are attributable to cell proliferation, immunohistochemical staining for Proliferating Cell Nuclear Antigen (PCNA) was performed (Fig. 2A) and demonstrated activation of cellular proliferation in the tendon bundle and the peritendinous tissues following injury, which peaked at 14 dpi. Furthermore, quantifying the percentage of PCNA^+^ cells at all timepoints after injury revealed increased cellular proliferation in the healing C57 tendons relative to the MRL tendons (Fig. 2B).

**Figure 2.**
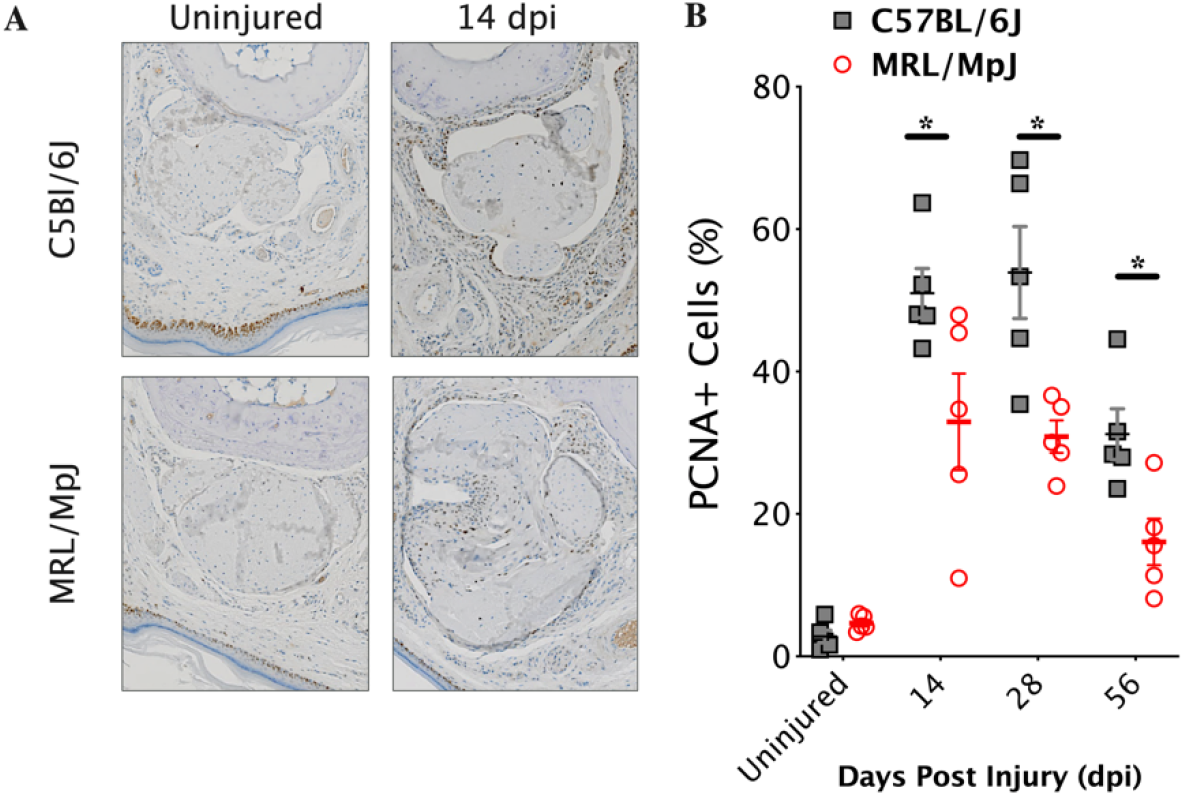
MRL flexor tendons heal with reduced cellular proliferation. (A) Representative images of proliferating cell nuclear antigen (PCNA) immunohistochemical (IHC) staining conducted before and 14 days post injury on 5 µm transverse paraffin sections and counterstained with hematoxylin, and then quantified (B) for percent PCNA-positive cells within the tendon and the surrounding peritendinous tissues (n=5).

The functional properties of injured tendons were evaluated using biomechanical testing, which involved displacement-controlled tensile stretching of the tendon to failure at 0.05□mm/sec. In terms of the structural properties, the injury resulted in significant decreases in peak tensile force and stiffness at 14 dpi in both the C57 and the MRL tendons. In contrast to the C57 tendons, which exhibited sustained reductions in peak tensile force and stiffness up to 56 dpi, the MRL tendons recovered uninjured levels of peak tensile force by 28 dpi (Fig. 3A) and were significantly stiffer (Fig. 3B) than the C57 tendons at all time points up to 56 dpi. Since the cross-section area of the uninjured MRL tendons are approximately 20% larger than the C57 tendons, we computed the corresponding stress-strain tensile curves and determined the effects of injury on the size-independent material properties. At 14 dpi, the tensile strength (peak stress) of the MRL was significantly reduced compared to uninjured tendon and the C57 injured tendon. The tensile strength of the C57 continued to decline at 28 dpi but recovered at 56 dpi. In contrast, the MRL tendon recovered the tensile strength to levels comparable to uninjured tendon as early as 28 dpi (Fig. 3C). Injury resulted in significant reductions in the elastic modulus of the C57 tendon, which did not improve over 56 dpi. While the elastic modulus of the uninjured MRL tendon was significantly reduced compared to the uninjured C57, injury resulted in reductions in the elastic modulus on 14, 28, and 56 dpi, which were not significantly different than uninjured tendon (Fig. 3D). Collectively, these data support the conclusion that MRL flexor tendons heal with reduced peritendinous adhesions and cellular proliferation and accelerated recovery of strength and stiffness.

**Figure 3.**
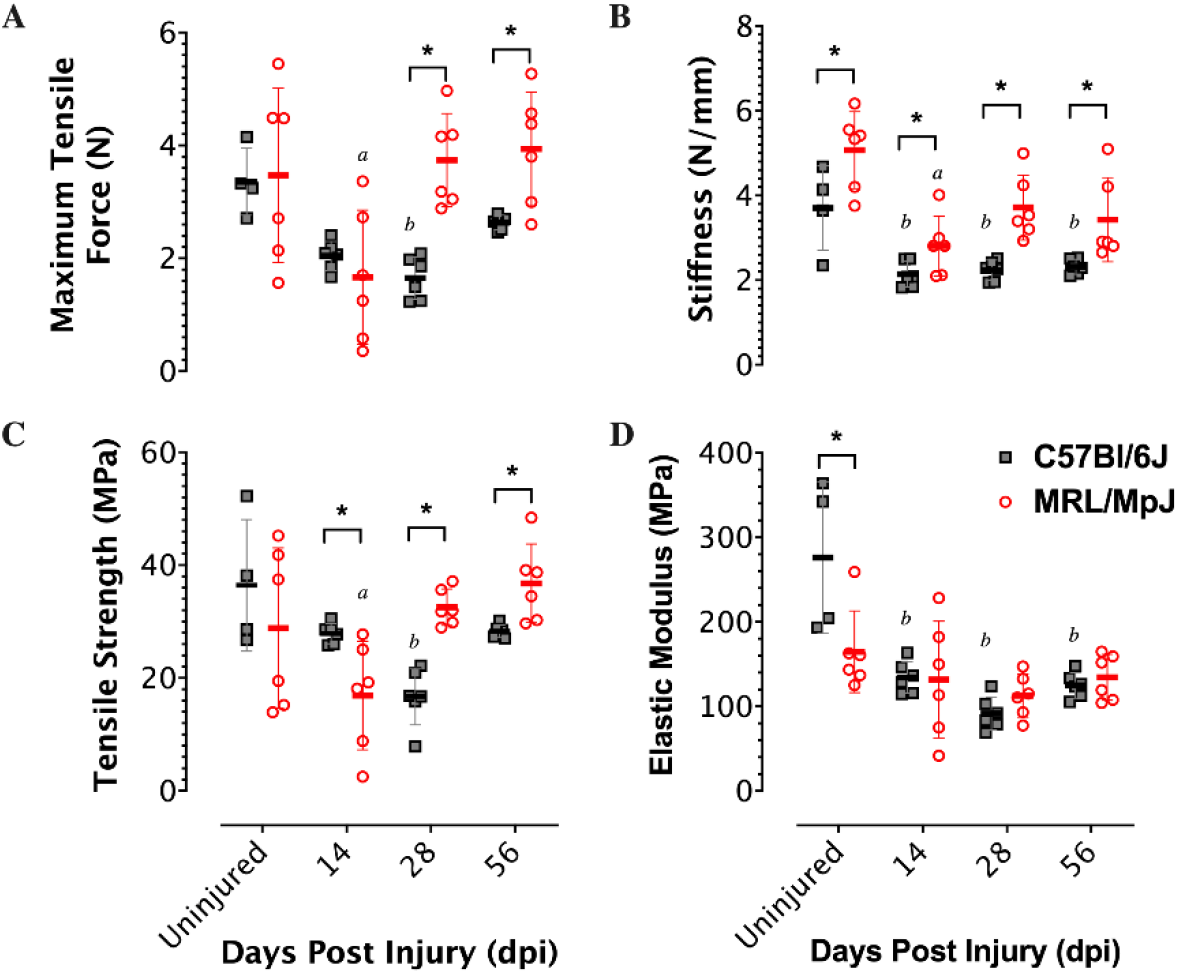
MRL flexor tendons heal with accelerated recovery of strength and stiffness (A) Maximum force, (B) strength, (C) stiffness. and (D) elastic modulus determined from displacement-controlled tensile testing of uninjured and injured flexor tendons of C57BL/6J and MRL mice (n=4-6). Asterisks indicate significant mouse strain differences (p < 0.05).

### Altered inflammatory, fibrotic, and cell cycle regulation in C57 and MRL tendon healing

To gain insights into the differences in the biological drivers of the tendon injury response between the C57 and MRL mice, we performed next generation sequencing of bulk tissue RNA (RNA-seq) at 7 days post-injury, which we reasoned to be a time point that defines the transition from the acute inflammatory phase to the reparative phases. Based on a 2-fold change (±2FC) and adjusted p-value < 0.05 differential expression criteria relative to healthy uninjured tendon, 891 genes were differentially expressed (594 upregulated and 297 downregulated) in the C57 tendons, and 1358 genes were differentially expressed (747 upregulated and 611 downregulated) in the MRL tendons. Of those, 406 genes (DEG) were differentially expressed, relative to respective uninjured controls, in both mice. We identified 22 significantly enriched Reactome pathways in the injured C57 tendon and 51 significantly enriched pathways in the injured MRL tendon based on False Discovery Rate (FDR) < 0.05. Functional annotation of these enriched gene sets identified remarkable differences in the transcriptional activity, wherein the gene sets enriched in the C57 injured tendon were functionally linked primarily to immune system signaling and extracellular matrix (ECM) organization (Fig. 4A), while the gene sets enriched in the MRL injured tendon were dominated by cell cycle pathways (Fig. 4B). Closer inspection of the top 15 Reactome pathways enriched in the C57 injured tendons (Fig. 5A) identified cytokine signaling, including interleukins 4, 10, and 13, and neutrophil degranulation as significantly enriched immune system pathways, and collagen formation (biosynthesis and assembly), ECM proteoglycans, ECM degradation, and integrin and non-integrin cell-ECM interactions as significantly enriched ECM organization pathways. In contrast, the top 15 Reactome pathways enriched in the MRL injured tendons (Fig. 5B) identified numerous cell cycle pathways related to mitotic phase, cell cycle checkpoints, and G1-S phase transition, as well as cell signaling by Rho GTPases and immune system signaling by interleukins and neutrophil degranulation.

**Figure 4.**
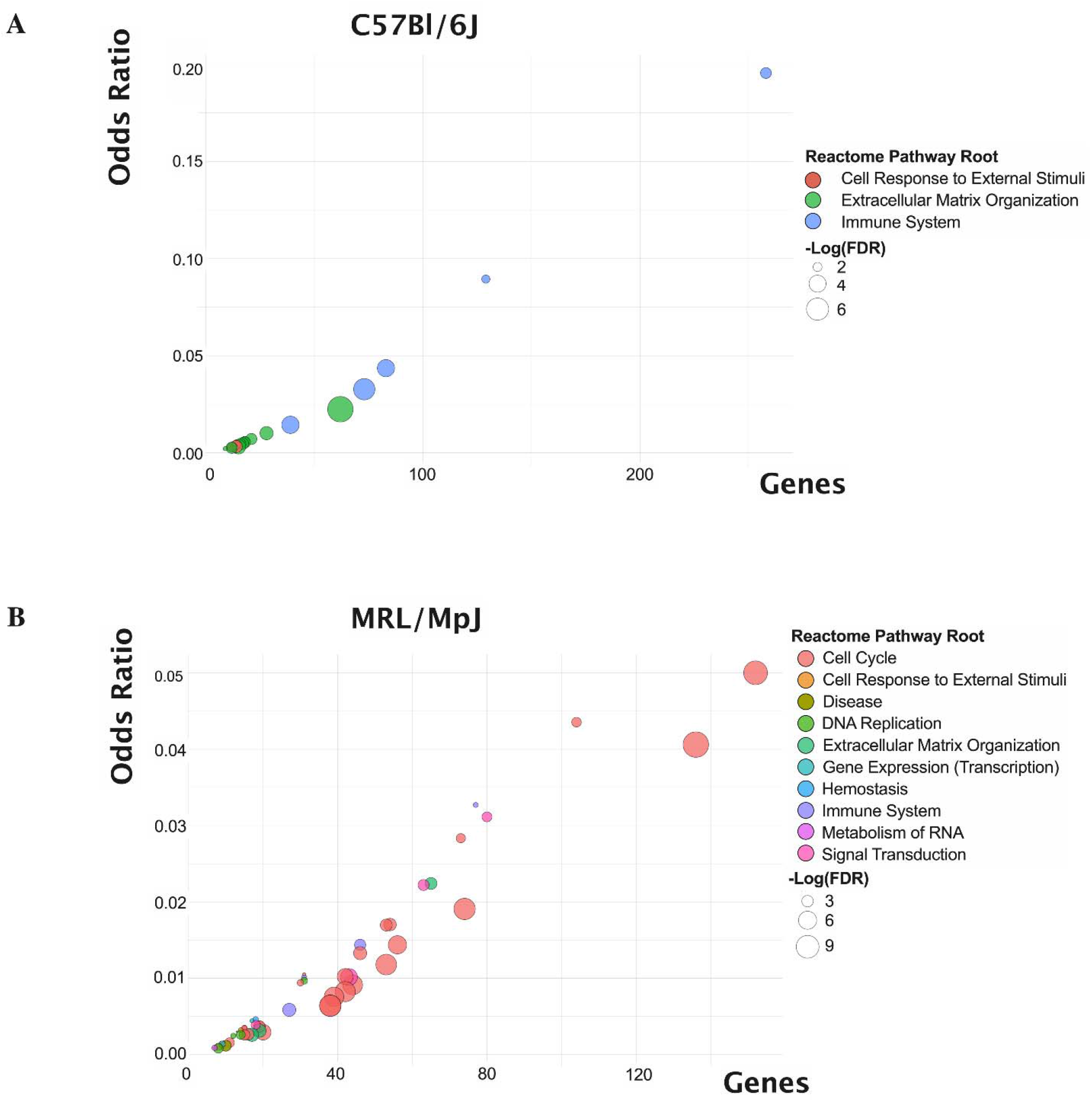
Functional annotation based on Reactome Pathways enrichment of 891 and 1358 differentially expressed genes (DEG) at 7 days post tendon injury in the C57 and MRL mice, respectively. Bubble plots of number of gene terms (x-axis) versus odds ratio (y-axis) represent 22 significantly enriched pathways in the injured C57 tendon (A) and 51 significantly enriched pathways in the injured MRL tendon (B) based on False Discovery Rate (FDR) < 0.05.

**Figure 5.**
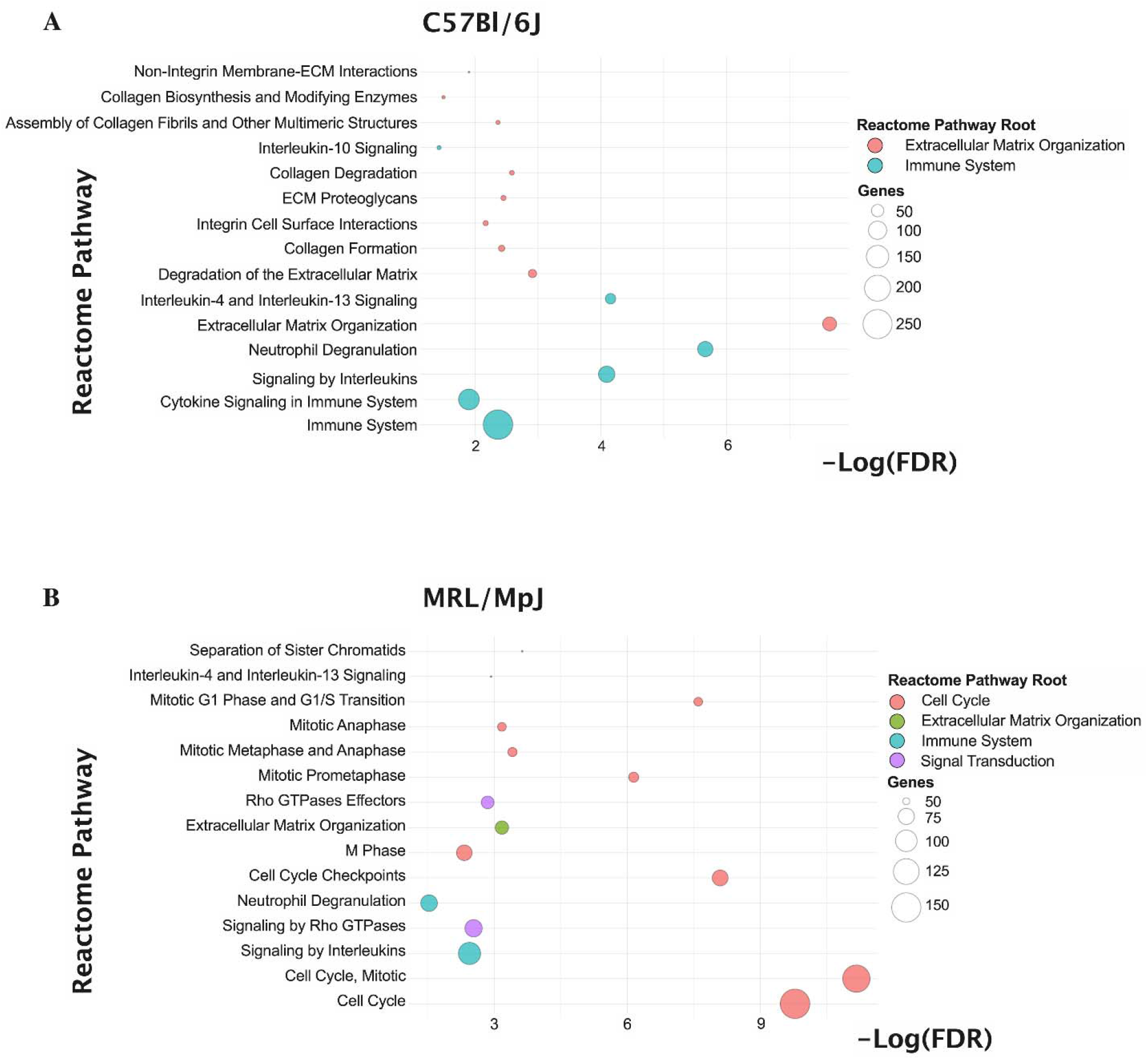
Bubble plots of the top 15 Reactome pathways enriched in the injured C57 tendon (A) and the MRL tendon (B), respectively. Bubble fill color represents the Reactome Pathway Root, while the size of the bubble represents -Log_10_(FDR) in A & B and the number of genes represented in the enriched pathway in C&D, respectively.

### Dichotomous roles for TGB1 in the C57 and MRL tendons

Ingenuity pathways analysis (IPA) of active or inhibited upstream transcriptional regulators were also examined to provide additional insights into the observed gene expression changes driving the altered mechanobiology of tendon healing. In the C57 injured tendons, the top 20 activated and inhibited upstream regulators (Fig. 6A) were associated with 184 and 62 unique genes, respectively, that enriched Gene Ontology (GO) terms related to chemotaxis and cell motility and migration, ECM organization, sprouting angiogenesis and blood vessel morphogenesis, positive regulation of cell growth and proliferation, inflammatory responses, collagen catabolism and regulation of phosphatidylinositol- and ERK1/2-signaling (Table S1). In the MRL injured tendon, the top 20 activated and inhibited upstream regulators (Fig. 6B) were associated with 325 and 304 unique genes, respectively, that enriched GO terms related to regulation of cell cycle, including mitotic cell cycle, sister chromatid cohesion, G1/S transition of mitotic cell cycle, mitotic cytokinesis, DNA replication, chromosome segregation, regulation of cell cycle, microtubule-based movement, metaphase plate congression, mitotic spindle organization, G2/M transition of mitotic cell cycle, negative regulation of cell cycle, and extracellular matrix organization (Table S2). Of particular note, was the inhibition of upstream regulators TP53, CDKN1A (p21), CDKN2A (p16), which are associated with DNA damage response and cell cycle arrest.

**Figure 6.**
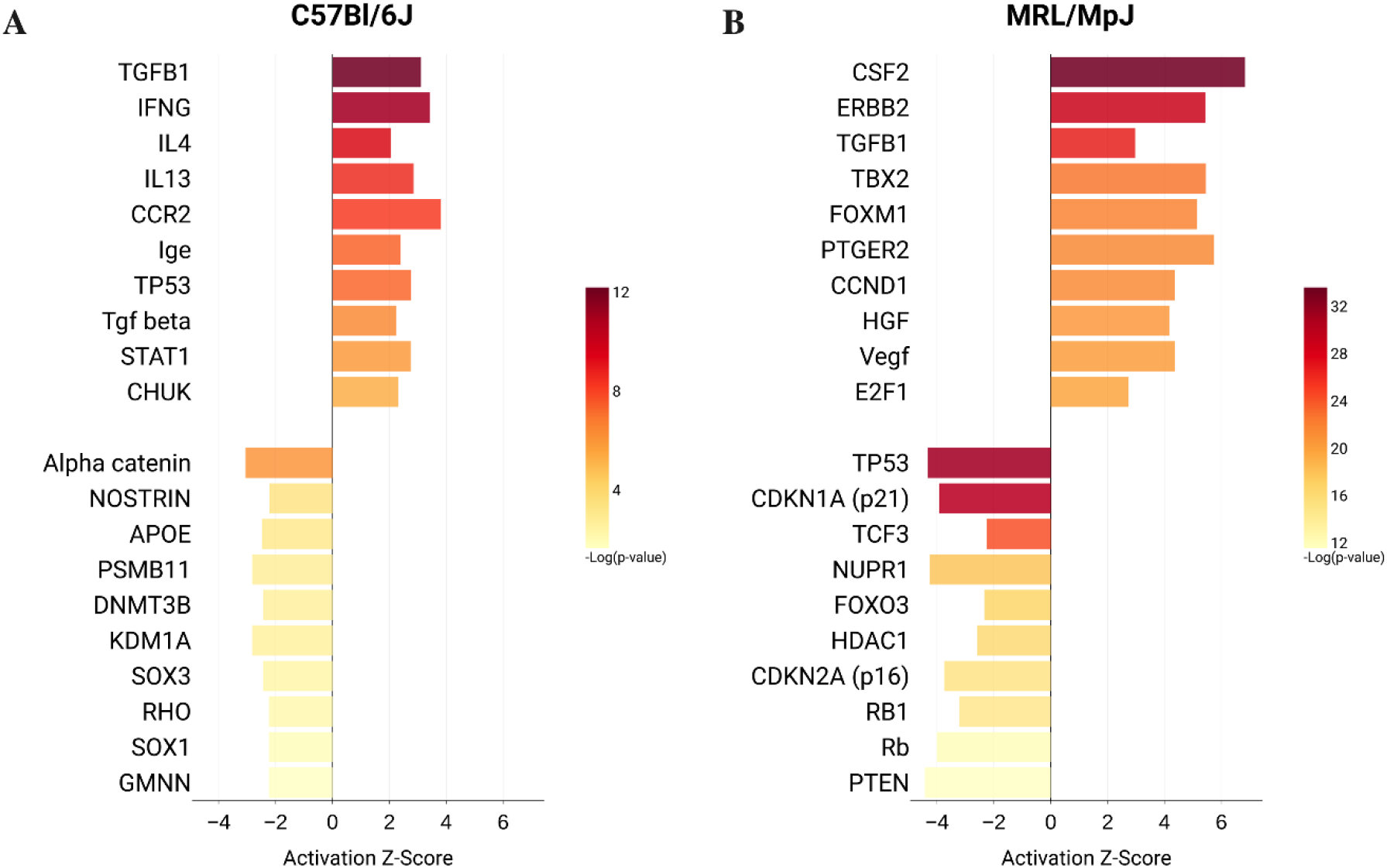
Ingenuity Pathway Analysis (IPA) Upstream Regulators during tendon healing in the C57 (A) and MRL (B) mice. Activation (Z-score ≥ 2, p-value of overlap < 0.05) or inhibition (Z-score ≤ −2, p-value of overlap < 0.05) was determined relative to uninjured tendon gene expression in either strain.

TGFB1 was a significantly activated Upstream Regulator in both the C57 (Activation Z-score = 3.11 and p-value = 6.4×10^−13^) and MRL (Activation Z-score = 2.97 and p-value = 1.1×10^−27^) injured tendons. GO functional annotation revealed that activation of TGF-β1 drives different biological pathways in the C57 and MRL injured tendons. In the C57 injured tendons, the 82 target genes of TGFB1 enriched GO pathways related to cytokine and chemokine signaling, ECM organization, and cell motility and migration (Fig. 7A). In contrast, 165 target genes of TGFB1 in the MRL enriched GO pathways associated with not only ECM organization and cytokine signaling but mostly with regulation of cell cycle, mitosis, and DNA binding and transcription (Fig. 7B).

**Figure 7.**
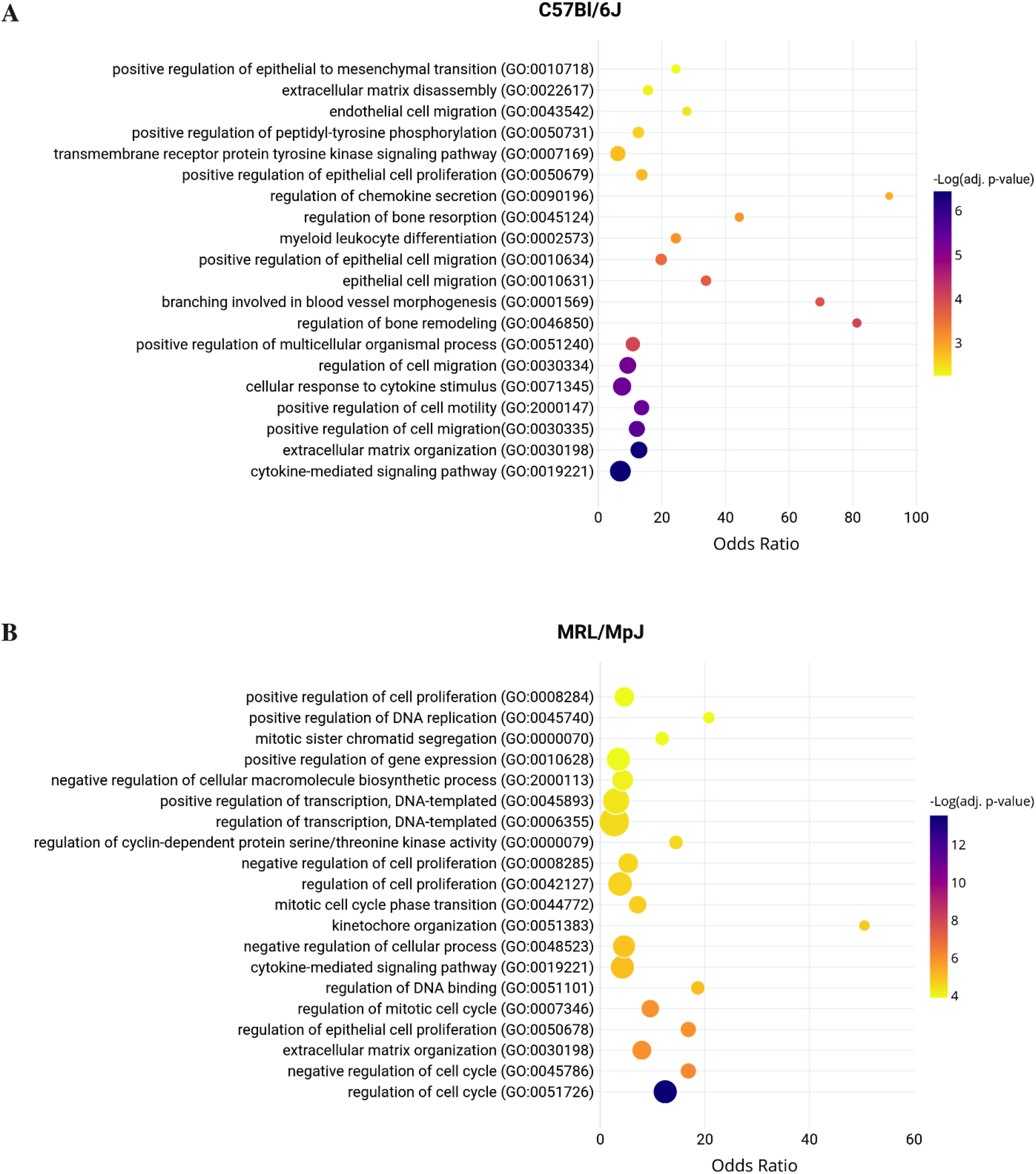
Biological Process Gene Ontology (GO) of 82 genes contributing to the activation of TGFB1 Upstream Regulator in the healing C57 tendon (A). Biological Process Gene Ontology (GO) of 165 genes contributing to the activation of TGFB1 Upstream Regulator in the healing MRL tendon (B). Bubble area corresponds to the number of genes in the enriched GO biological process.

### Reduced circulating cytokines and chemokines in the MRL injured mice

In order to evaluate the roles of the immune response and systemic inflammatory signaling on the tendon healing, sera were extracted from whole blood draws from mice at different time points up to 14 dpi and analyzed for senescence associated secretory phenotype (SASP) proteins including inflammatory cytokines/chemokines, TGF-βs, and MMPs. In general, C57 and MRL mice exhibit similar circulatory concentrations of serum proteins in their uninjured state (Fig. 8A). Upon injury, the temporal changes in the circulating proteins in the C57 mice were in general more pronounced than in the MRL mice. Analyzed proteins were clustered in 4 major classes based on Pearson’s correlations (Fig. 8B). In the first cluster, the chemokine CXCL9 (MIG) showed no significant difference over time in the C57 but increased significantly over time in the MRL injured tendons. Cluster 2 proteins showed no significant differences over time or between C57 and MRL mice, which included TGF-βs (1,2,3), MMP-3 and CCL 4 (MIP-1β). Cluster 3 proteins, which included IL-1α, -12 (P40), -13, MMP-2 and -8, CSF1 (M-CSF) and 3, CCL5 (RANTES) and 11 (Eotaxin), CXCL2 (MIP-2) and 5 (LIX), were significantly higher in the C57 mice compared to the MRL mice, but did not vary significantly over time. Cluster 4 proteins showed significant changes over time in the C57 mice only, peaking at either 1 dpi (IL-2 and -9, MMP-12, and CCL3 (MIP-1α)) or 5 dpi (IL-1β, -2, -4, -5, -6, -7, 10, -12 (P70), -17, TNF-α, LIF, MMP-9, CCL2 (MCP-1) and CXCL1(KC) and 10 (IP-10)). In contrast circulation levels of these proteins were significantly lower and remained mostly unchanged over time in the MRL mice. Protein association networks of cluster 3 (Fig. 8C) and cluster 4 (Fig. 8D) circulating proteins visualized in String DB identified strong functional associations, which involved cytokine signaling in immune system and more specifically signaling by interleukins, chemokine signaling and activation of matrix metalloproteinases.

**Figure 8.**
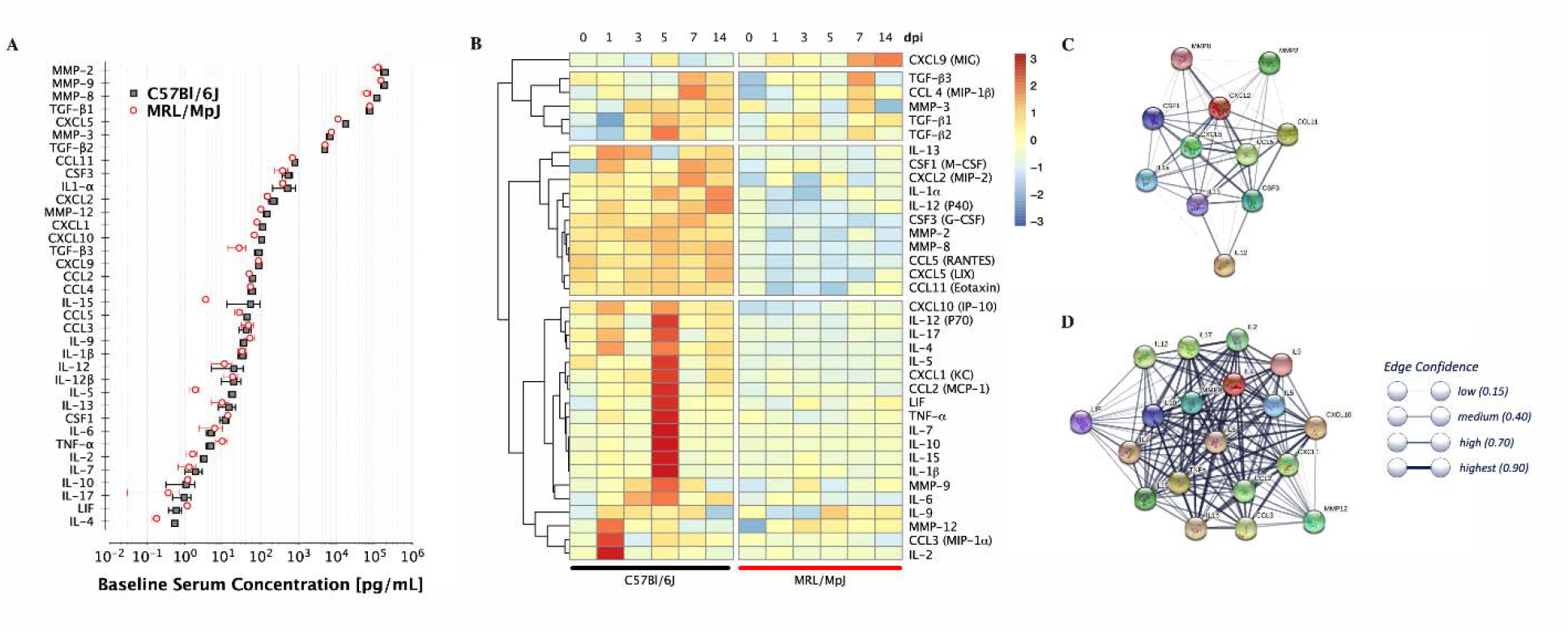
Serum protein analysis indicates temporal systemic upregulation of several circulating enzymes, cytokines, and chemokines in the C57 mice following tendon injury, but not in the MRL mice. (A) Baseline concentrations of serum proteins pre-injury plotted in order of concentration shows no significant differences between C57 and MRL mice. (B) Heatmaps for circulating serum enzymes, cytokines, and chemokines concentration (normalized to day 0 C57 controls) changes over time post flexor tendon injury (dark red indicates > +3 fold increase and dark blue indicated > -3 fold decrease compared to preinjury serum concentration). Dendrogram represent clustering based on Pearson’s correlations. Each box represents the average of 3 mice at each time point. String DB representation of physical and functional association network of cluster 3 (C) and cluster 4 (D) proteins.

## Discussion

Development of fibrotic flexor tendon adhesions, particularly in zone II of the hand (Elliot et al., 2016, Tang et al., 2013), limits their natural gliding function, strength, and intricate joint flexion (Tang, 2013, Tang et al., 2013, Zhou et al., 2017). Recent biological advances in scarless repair of tendon injuries (George et al., 2020, Paredes et al., 2020, Paredes et al., 2018, Freeberg et al., 2019) have begun to elucidate the complex cellular and transcriptional interplay in restoring native tissue properties to injured tendons. For example, previous studies demonstrate that MRL tendon injuries heal to near normal biomechanical properties (Lalley et al., 2015, Arble et al., 2016), have reduced infiltrating immune cells (Lalley et al., 2015), attenuated inflammatory expression (George et al., 2020, Paredes et al., 2018), lower catabolic enzyme activity (Sebastian et al., 2018), heightened MMP activity (George et al., 2020), and improved ECM alignment (Paredes et al., 2020). These reports are consistent with our current findings in flexor tendons, but the biological drivers of this regenerative tendon healing phenotype have hitherto not been fully understood.

The objective of our study was to uncover the mechanisms of improved healing of MRL flexor tendon injuries to understand better the limitations of natural healing in common C57 mouse tendons. We first investigated the longitudinal histomorphometry and biomechanical properties of MRL and C57 mice flexor tendons after a zone II partial laceration injury. We demonstrated that the MRL flexor tendons heal with reduced tenocyte cellularity, reduced peritendinous adhesions, and accelerated recovery of their biomechanical properties of tensile strength, stiffness, and elastic moduli. We then performed bulk tissue transcriptomic analysis using next-generation RNA sequencing (RNA-Seq) to characterize enriched transcriptional pathways in the early reparative phase. These experiments uncovered dichotomous roles for TGFB1 in the healing of C57 and MRL tendons. Specifically, we identified TGFB1 as a major driver of positive enrichment of cytokine signaling, neutrophil degranulation, and ECM degradation pathways in injured C57 flexor tendons, and, by contrast, negative enrichment of cell cycle regulation and senescence pathways in the injured MRL flexor tendons. To validate these transcriptional observations, we investigated the systemic inflammatory response by analyzing the cytokines, chemokines, and growth factors in sera extracted from peripheral blood at different phases of healing. These assays revealed significantly lower circulatory concentrations of SASP proteins in the MRL relative to the C57 despite comparable baseline (day 0) serum concentrations. These findings provide insights into novel molecular pathways in tendon healing that could be therapeutic targets for regenerative tendon repair.

TGF-β is critical to the modulation of tendon healing and the effects of its isoforms are quite distinct. In flexor tendons, TGF-β1 expression is auto-inductive and is initially produced by inflammatory cells (Wojciak and Crossan, 1994, Wojciak and Crossan, 1993). TGF-β1 stimulates fibroblast proliferation and migration, synthesis of collagen and fibronectin (Wojciak and Crossan, 1994, Klein et al., 2002), inactivation of MMPs (Wang and Hirschberg, 2003) and myofibroblast differentiation (Kaji et al., 2020). Chang et al. reported increased expression of TGF-β1 in both tenocytes and infiltrating fibroblasts and inflammatory cells from the tendon sheath in a rabbit zone II flexor tendon healing model (Chang et al., 1997). The inflammatory cells stimulate synovial and epitenon fibroblasts, partly through TGF-β1, to produce fibronectin, a marker of scarring (Wojciak and Crossan, 1994, Wojciak and Crossan, 1993). The role of TGF-β1 in scar formation is accentuated by the observation that while fetal cutaneous wound healing and tendon repair are scar-free (Beredjiklian et al., 2003, Krummel et al., 1988), the *in utero* administration of TGF-β1 to fetal rabbit wounds results in fibrotic scarring similar to that observed in adult rabbits (Krummel et al., 1988). While there are numerous differences between adult and embryonic wound healing, the temporal expression of the TGF-β isoforms are highly associated with scar formation or the lack thereof, respectively. For example, TGF-β1 and TGF-β2 are abundant early on in adult wounds, but they are very low and nearly absent in embryonic scar-less wounds (Ferguson and O’Kane, 2004, Kuo et al., 2008). TGF-β3, which is highly induced in developing skin and embryonic scar-less wounds, is induced late and at low levels in adult wounds only after TGF-β1 levels start to fall (Ferguson and O’Kane, 2004). The temporal expression of TGF-β isoforms in healing tendons have been less extensively studied *in vivo* (Chan et al., 2008, Tsubone et al., 2004). The sparse data suggest that TGF-β1 levels peak as early as 4 days and persist for as long as 28 days in the injured adult rat patellar tendon *in vivo*. In contrast, TGF-β2 and TGF-β3 levels increase after 7 days as TGF-β1 levels begin to slowly decrease (Chan et al., 2008).

In our study, we did not observe changes in circulating TGF-β1, 2, or 3 levels over time or between the C57 and MRL mice. However, we did observe dichotomous downstream effects of TGFB1 gene regulatory networks between the C57 and MRL healing tendons locally. In particular, genes regulated by TGFB1 in the C57 tendons positively enriched pathways relevant to cytokine signaling in chemotaxis and inflammation, ECM organization, angiogenesis, and epithelial to mesenchymal transition, whereas the TGFB1 regulated genes in the MRL tendons negatively enriched pathways predominately related to cell cycle regulation and senescence. These results corroborate reports of dual roles for TGF-β in different contexts, such as being protective and cytostatic against tumorigenesis but also a promoter of tumor progression, invasion, and metastasis in established tumors (Lebrun, 2012). Dual roles of TGF-β have also been reported in autoimmune-induced tissue damage, wherein reduced TGF-β signaling in immune cells can lead to inflammatory dysregulation triggering the production of TGF-β in target organs, which subsequently leads to fibrogenesis and organ failure (Saxena et al., 2008). It is possible that the effects of TGF-β in the injured tendon are intertwined with the inflammatory cytokines and chemokines in the site of injury and systemically. It is also possible that intrinsic genetic differences in the tendon fibroblasts and infiltrating inflammatory cells could explain the dichotomous responses downstream of TGFB1. A third possibility is that the innate tissue properties in the MRL mouse tendon (Paredes et al., 2020) can modulate the effects of TGF-β1 resulting in accelerated repair. These possibilities warrant further investigation in future studies.

RNA-Seq analyses revealed negative enrichment of cell cycle regulation in the MRL tendons compared to the C57 healing flexor tendons who exhibited positive enrichment of inflammatory and ECM organization pathways, driven differentially by TGFB1 regulated gene sets. The increased responsiveness of injured C57 tendons to circulating inflammatory cytokines, chemokines, and growth factors may be associated with its positive enrichment of cell cycle evidenced by cyclin D1 (*ccnd1*) pathways. In contrast, injured MRL tendons negatively enriched pathways associated with cell cycle regulators TP53, CDKN1A (p21), CDKN2A (p16), RB1, FOXO3, and PTEN. The enrichment of these cell cycle regulators is associated with cellular senescence(Campisi and d’Adda di Fagagna, 2007, Tchkonia et al., 2013). Cell cycle regulation and cellular senescence have recently become therapeutic targets in a number of conditions including fibrosis. Specifically, the cell cycle regulator p53 is induced in multiple fibrotic conditions such as idiopathic pulmonary fibrosis (IPF) and kidney fibrosis. In the absence of p53, bleomycin induced pulmonary fibrosis is prevented(Shetty et al., 2017) and its absence reduces chronic renal injury(Higgins et al., 2018). Furthermore, the antifibrotic agent used to treat IPF, Pirfenidone, has been shown to induce cell cycle arrest (Ruwanpura et al., 2020, Sun et al., 2018, Usugi et al., 2019). Reduced cell apoptosis, cell cycle regulation through p21 (*cdkn1a*)(Jablonski et al., 2019, Bedelbaeva et al., 2010), p53 (*trp53*), and p16 (*cdkn2a*), increased cell proliferation(Bedelbaeva et al., 2010), and enhanced stem cell function(Naviaux et al., 2009) may command augmented regenerative capacity. Given that PTEN, a tumor suppressor gene, is a natural inhibitor of the phosphoinositide 3-kinase (PI3K) and of the mammalian target of rapamycin (mTOR) pathway, the negative enrichment of PTEN regulated genes provide a mechanistic link between mTOR-regulated cell cycle and scar healing in tendon. Reduced PTEN signaling has been associated with renal fibrosis via p53-, smad3-, and PI3K/Akt/mTORC1-dependent induction of cellular senescence (Samarakoon et al., 2015). A recent genome-wide association study (GWAS) of susceptibility to idiopathic pulmonary fibrosis strongly implicated mTOR signaling as a causal driver of fibrosis pathobiology in humans (Allen et al., 2020). Interestingly, there are two currently active clinical trials investigating the safety, tolerability and pharmacokinetics (NCT03502902) and the efficacy (NCT01462006) of mTOR inhibitors as disease modifying drugs for IPF. Collectively, these reports provide evidence linking TGF-β signaling to PTEN and mTOR-induced cellular senescence as a mediator of fibrosis.

While the prolonged systemic response in the C57 is surprising given the focal nature of the injury, the diminished response in the MRL is likely associated with decreased recruitment of inflammatory cells to the injury site similar to the scar-free healing of fetal wounds(Ferguson and O’Kane, 2004). Interestingly, chemokine (C-X-C motif) ligand 9, CXCL9 is the single cytokine in cluster 1 and is transiently upregulated in MRL healing. In other fibrotic conditions increased CXCL9 has been beneficial to patient function and reduced fibrosis(Nukui et al., 2019, Sahin et al., 2012). A study of serum cytokine levels of patients with hypersensitivity pneumonitis pulmonary fibrosis demonstrated a worsening lung function with reduced levels of CXCL9(Nukui et al., 2019). Additionally, systemic administration of CXCL9 prevented the development of CCL4 induced liver fibrosis in a murine model(Sahin et al., 2012). Altogether, the observed elevation of CXCL9 in the MRL sera after injury points to a potential therapeutic benefit of CXCL9 administration, which merits testing in tendon injury. In addition, proteins in clusters 3 and 4 can potentially be viewed as serum biomarkers or potentially candidates for therapeutics inhibition to mitigate fibrovascular scar healing in tendon.

This study has yielded novel insights into the transcriptional upstream regulators that drive enhanced tendon healing and fibrosis. However, there are limitations that qualify the conclusions of our study. First, our study was limited in assessing transcriptional regulation in bulk tendon tissue in native tendon and 7 days after injury. Whole-genome transcriptional profiling from the injured tendon tissue site represents a heterogeneous cell population of tenocytes, fibroblasts, myofibroblasts and immune cells, which could be resolved by single-cell RNA-sequencing in the tendon as recently described(De Micheli et al., 2020, Kelly et al., 2020). Future analysis will benefit from the utilization of single cell RNA-sequencing or spatial transcriptomics to elucidate signaling mechanisms at a cellular level given the involvement of the heterogeneous cellular populations at the tendon injury site and at later time-points that would represent the fibro-proliferative and remodeling phases of wound healing.

In summary, this study identified critical cell cycle regulation and differentiation roles for TGFB1 signaling that contribute to improved tendon repair following injury in MRL flexor tendons but not in the C57 tendons. Uncovering the unique cellular and molecular pathways in which TGFB1 induces regenerative tendon healing in the MRL model could help identify therapeutic targets for regenerative tendon healing.

## Materials and Methods

### Animal Care and Flexor Tendon Surgery

All animal procedures were conducted in compliance with protocols approved by the University of Rochester Committee on Animal Research (UCAR). The mouse strains utilized in this study were the C57BL/6J (C57) and the Murphy Roths Large MRL/MpJ (MRL) mice. Breeding pairs for the C57BL/6J (stock number: 000664) and MRL/MpJ (stock number: 000486) were obtained from the Jackson Laboratory (Bar Harbor, ME) and bred in-house. Nine to eleven-week-old male mice from each strain were randomized into experimental groups. The mouse surgery protocol involves a partial laceration of the deep digital flexor (DDF) tendon of the 3rd digit in the hind paw, as previously described(Freeberg et al., 2018). Surgeries were performed under a stereomicroscope on the hind paws using sterile aseptic techniques. Mice were anesthetized with 60 mg/kg ketamine and 4 mg/kg xylazine. A transverse incision was made on the 3rd digit between the metatarsophalangeal (MTP) and the proximal interphalangeal (PIP) joints. A transverse, mediolateral cut across roughly 50% of the DDF tendon width was completed. The skin incision was closed using interrupted sutures (Ethicon Suture, V950G, 9-0). To minimize pain, all mice were subcutaneously administered Buprenorphine at a dose of 0.05 mg/kg every day for up to 72 hours. Blood was collected from mice whose hind paws were used for gene expression analysis.

### Biomechanical Testing

Tensile elastic biomechanical testing was completed as previously described(Freeberg et al., 2018). The DDFT was released at the myotendinous junction, precisely dissected along the tibia and tarsal bones, and released from the tarsal tunnel pulleys. The middle (3rd) tendon was isolated, with the tendon-to-bone attachment (enthesis) left intact for mounting and gripping. Tendons were loaded in a custom uniaxial microtester (eXpert 4000 MicroTester, ADMET, Inc., Norwood, MA), and gripped between two stainless steel plates with the 3^rd^ digit tip superglued between. After a preloading protocol to 0.05 N, tendon width, thickness, and gauge length were measured while the tissue was immersed in a 1x PBS bath and imaged on the microscope. Cross sectional area was assumed to be an ellipse. Based on each sample’s total gauge length, each sample underwent uniaxial displacement-controlled biomechanical testing as follows; 1% strain preconditioning for 10-cycles, returned to initial gauge length to rest for 5 minutes, stretched to 5% strain (0.05 strain/8 seconds) and held for 10 minutes to allow for stress-relaxation, returned to initial gauge length to rest for 5minutes, and concluded with a displacement-controlled ramp to failure at 0.05 mm/sec. Structural and material properties were determined from the ramp to failure force-displacement and stress-strain curves, respectively. All mechanical testing analysis was completed with a custom Matlab (MathWorks, Natick, MA) code. Sample size of N=6 for each mouse strain and each experimental timepoint.

### Histology and Immunohistochemistry

The hind paw was transected near the tarsal bones and prepared for histology by scoring the skin superficial to the footpad to enhance 10% Neutral Buffered Formalin (NBF) tissue infiltration. Tissues were fixed in 10% NBF for 48hrs at room temperature (RT), followed by decalcification in calis-EDTA (pH 7.2-7.4) for 7 days at RT. The tissues were then dehydrated in an ethanol gradient and embedded in paraffin. Paraffin-embedded tissues were cut axially into 5μm thick sections between the metatarsophalangeal (MTP) joint and the A3 pulley of each sample, then stained for 15 seconds with Mayer’s Hematoxylin. Sections for histomorphometric analysis were immediately cover slipped with a water-based mounting medium (Faramount Aqueous Mounting Medium, Dako, #S3025), whereas sections for immunohistochemistry (IHC) were cover slipped after IHC.

For immunohistochemistry, paraffin-embedded and sectioned tissues were collected at 0, 14, 28, and 56 days post injury (dpi). Slides were incubated in citrate buffer (10mM Na citrate, pH 7.2) for 3hrs at 70°C for antigen retrieval, circled with a hydrophobic barrier pen (Vector Laboratories, Burlingame, CA), and blocked with an endogenous peroxidase/ alkaline phosphatase inhibitor (Bloxall, Vector Laboratories). After washing with PBST (PBS + 0.1% Tween-20), they were incubated for 1hr at room temperature in blocking solution (5% normal horse serum + PBST). Sections were incubated for 1hr at RT with the primary antibody mouse monoclonal PCNA (ab29, Abcam) at 1:1000 (1μg/mL) dilution. Followed by Vectastain Elite Kit with DAB chromogen (Vector Laboratories). Immunohistochemical sections were rinsed with deionized water, counterstained with Mayer’s Hematoxylin, cover slipped with a water-based mounting medium (Faramount Aqueous Mounting Medium, Dako, #S3025), and imaged with a VS120 Virtual Slide Microscope (Olympus, Waltham, MA).

### Quantification of Histomorphometric and Immunohistochemical Images

For histomorphometry, paraffin-embedded tissues were prepared as described above. Brightfield images were obtained from the slide scanner and processed using Visiopharm image analysis software v.6.7.0.2590 (Visiopharm, Hørsholm, Denmark). Regions of interest were manually drawn to define only the tendon tissue area inside the synovial sheath. Using a custom Visiopharm program, imaged sections were Bayesian classified and thresholded to automatically train the program to specific pixel RGB colors and intensities. To improve accuracy and repeatability, post-processing analyses defined cell nuclei area >2µm^2^, synovial sheath space (SS) 5µm^2^ < SS < 5000µm^2^, eroded local minima pixel intensities for each colorimetric label, and labeled blank space within the tendon body as intratendinous space. Magnified brightfield images of hematoxylin stained sections (10X, 20X) were manually delineated to distinguish the superficial digital flexor tendons (SDFT) from the deep digital flexor tendons (DDFT). The following tissue measurements were taken: DDF tendon cross-sectional area, synovial sheath space (SS), tendon extracellular matrix (ECM), intratendinous cell nuclei count, intratendinous cellular density, and length of DDFT perimeter adhered to subcutaneous tissues(Freeberg et al., 2018). Each histomorphometric outcome measure was quantified using Visiopharm software averaged over 4-5 animals per experimental group.

### RNA Extraction

Tendon tissue was collected from the injury site for RNA extraction and subsequent gene expression analysis by the URMC Genomics Core. Briefly, partially lacerated or uninjured tendon tissue was flash frozen in liquid nitrogen and stored at -80°C until time of extraction. Total RNA was isolated from single tendons utilizing Trizol (ThermoFisher, #15596026) extraction methods and a bullet blender for tissue homogenization. RNA concentration was determined with the NanoDrop 1000 spectrophotometer (NanoDrop, Wilmington, DE) and RNA quality assessed with the Agilent Bioanalyzer 2100 (Agilent, Santa Clara, CA).

### Next-Generation Sequencing (NGS) Data Processing and Alignment

Samples of 1ng of total RNA were pre-amplified with the SMARTer Ultra Low Input kit v4 (Clontech, Mountain View, CA) per manufacturer’s recommendations. The quantity and quality of the subsequent cDNA was determined using the Qubit Flourometer (Life Technnologies, Carlsbad, CA) and the Agilent Bioanalyzer 2100 (Agilent, Santa Clara, CA). 150pg of cDNA was used to generate Illumina compatible sequencing libraries with the NexteraXT library preparation kit (Illumina, San Diego, CA) per manufacturer’s protocols. The amplified libraries were hybridized to the Illumina single end flow cell and amplified using the cBot (Illumina, San Diego, CA). Single end reads of 100nt were generated for each sample. Raw reads generated from the Illumina HiSeq. 2500 sequencer were demultiplexed using bcl2fastq version 2.19.0. Quality filtering and adapter removal are performed using Trimmomatic version 0.36 with the following parameters: “TRAILING:13 LEADING:13 ILLUMINACLIP:adapters.fasta:2:30:10 SLIDINGWINDOW:4:20 MINLEN:15”. Quality reads were then mapped to the Mus musculus reference sequence (GRCm38.p5) with STAR_2.5.2b with the following parameters: “□□twopassMode Basic □□runMode alignReads □□genomeDir ${GENOME} □□readFilesIn ${SAMPLE} □□outSAMtype BAM SortedByCoordinate □□outSAMstrandField intronMotif □□outFilterIntronMotifs RemoveNoncanonical”. Differential expression analysis and data normalization were performed using default methods in DESeq. 2□1.14.1 with an adjusted *p* value threshold of 0.05.

### Bioinformatics and Statistical Analyses of DEG

Differential expression analysis and data normalization were performed using DESeq. 2□1.14.1 on RNA extracted from injured tendons at 7 days post injury (dpi) relative to uninjured expression levels within each strain independently. DEG was identified by filtering DeSeq. 2 pairwise comparisons for biological (ABS(Log_2_FC) > 1) and statistical significance (adjusted *p* < 0.05). Functional enrichment of upstream regulators was completed in pathway enrichment analysis using Ingenuity Pathway Analysis (IPA; http://www.ingenuity.com). Default settings were used to determine upstream regulators using the global molecular network contained in the IPA knowledge base. Downstream targets of the upstream regulator TGF-β1 identified in IPA were exported for further functional analysis using Enrichr (https://maayanlab.cloud/Enrichr/) and Reactome (https://reactome.org).

### Data Availability

The RNA□seq raw and processed data were deposited in the Gene Expression Omnibus under accession GEO: XXXXX.

### Multiplex Serum Protein Assay

Peripheral blood was collected via cardiac puncture from uninjured and at 1, 3, 5, 7, and 14 days post injury (dpi). The collection tube was allowed to clot completely at room temperature then centrifuged to isolate blood serum from coagulate. Blood serum was transferred to cryovials for storage at -80°C before shipping to EVE Technologies for multiplex protein analysis (EVE Technologies 3415A – 3 Ave., N.W. Calgary, AB Canada T2N 0M4). Serum protein analysis was performed with three analytical arrays of cytokines, chemokines, and transforming growth factor beta isoforms. Sample size was 3 mice per strain per time-point. Serum protein concentrations for Day 5 C57 contain only two samples as the third serum sample was determined to be a statistical outlier.

### Statistical Analyses

All histomorphometry and biomechanics data were graphed and statistically analyzed with GraphPad Prism (GraphPad Software). Significant differences (p < 0.05) for all data were determined using a 2-way ANOVA and Bonferroni-corrected multiple comparison post-tests. Differential expressional analysis and plots was generated in R Studio with an adjusted *p*-value threshold of 0.05.

## Acknowledgements

The study was supported by grant numbers R01AR056696, R01AR073169, UG3TR003281, and P30AR069655 from NIAMS/NIH. MTF was supported by the Training In Orthopaedic Research Program funded by NIH T32AR053459. The content is solely the responsibility of the authors and does not necessarily represent the official views of the National Institute of General Medical Sciences or NIH. We thank Brittany Strauss, Sarah Mack and Kathleen Maltby for histology assistance.

## Author Contributions

Conceptualization: HA, DA, JGK, MTF, AvW. Methodology: JGK, DA, MTF, JA, MB, AL. Software: JGK, MTF, JM. Formal Analysis: JGK, MTF, DA, JA, JM, AVW, HA. Investigation: JGK, DA, MTF. Writing – Original Draft: JGK, MTF, HA. Writing – Reviewing and Editing: JGK, MTF, HA, AL, MB, AvW. Visualization: JGK, MTF, HA. Supervision: HA. Project Administration: HA. Funding Acquisition: HA.

